# *otpb* functions in a Lef1-dependent transcriptional network required for expression of the stress response inhibitor *crhbp* in the zebrafish hypothalamus

**DOI:** 10.1101/2023.06.07.544119

**Authors:** Priscilla Figueroa, Jennifer Cheng, Guangning Wang, Cade Kartchner, Debora Brito de Andrade, Harrison Watters, Ethan Crispell, Richard I. Dorsky

**Affiliations:** Department of Neurobiology, University of Utah, Salt Lake City, UT 84112, USA

**Author notes:** Funding Sponsor: NIH (NINDS), Grant # R01NS082645.

## Abstract

The vertebrate hypothalamus regulates physiological and behavioral responses to environmental stimuli through the function of evolutionarily-conserved neuronal subpopulations. Our previous work found that mutation of zebrafish *lef1*, which encodes a transcriptional mediator of the Wnt signaling pathway, leads to the loss of hypothalamic neurons and behavioral phenotypes that are both associated with stress-related human mood disorders However, the specific Lef1 target genes that link neurogenesis to behavior remain unknown. One candidate is *otpb*, which encodes a transcription factor with known roles in hypothalamic development. Here we show that *otpb* expression in the posterior hypothalamus is Lef1-dependent, and that like *lef1*, its function is required for the generation of *crhbp*+ neurons in this region. Transgenic reporter analysis of a *crhbp* conserved noncoding element suggests that *otpb* participates in a transcriptional regulatory network with other Lef1 targets. Finally, consistent with a role for *crhbp* in inhibiting the stress response, zebrafish *otpb* mutants exhibit decreased exploration in a novel tank diving assay. Together our findings suggest a potential evolutionarily-conserved mechanism for the regulation of innate stress response behaviors through Lef1-mediated hypothalamic neurogenesis.

## Introduction

The hypothalamus plays a fundamental role in maintaining physiological homeostasis in response to changing environmental conditions. Signals from hypothalamic neurons, including neuropeptides and hormones, function through the nervous and endocrine systems to activate responses to external stimuli, and also to inhibit those responses through negative feedback signaling. Understanding the genetic and molecular mechanisms that regulate the development of hypothalamic neurons is important due to these critical homeostatic functions, and the association of hypothalamic dysfunction with diseases and behavioral disorders. Recent work has identified genes required for the differentiation of some hypothalamic neuronal subtypes, as well as transcription factors that regulate their expression of specific neuropeptides and hormone signals (Xie and Dorsky, 2017). However, the mechanisms underlying the development of other subtypes that control fundamental neuronal and behavioral responses remain unknown.

We previously identified required roles for Lef1, a transcriptional mediator of the Wnt signaling pathway, in regulating hypothalamic neurogenesis and innate behavior (Xie et al., 2017) Zebrafish *lef1* loss-of-function mutants fail to generate multiple neuron subtypes in the posterior recess (PR) of the caudal hypothalamus, including neurons that express *crhbp*, which encodes an inhibitor of corticotropin releasing hormone (Crh) (Li et al., 2016; Seasholtz et al., 2002). Crh signaling plays a primary role in mediating the physiological and behavioral responses to stress (Vom Berg-Maurer et al., 2016), and mouse *Crhbp* knockouts exhibit stress response-related behavioral phenotypes consistent with those observed in zebrafish *lef1* mutants, including decreased exploration of a novel environment (Karolyi et al., 1999). This role of Crh signaling extends to invertebrates including *Drosophila* (Mohammad et al., 2016), in which we found that the *Lef1* ortholog *pangolin* is required for expression of a *crhbp* ortholog in the developing neuroendocrine brain region (Xie et al., 2017). Together, these data suggest that Lef1-dependent generation of *crhbp*-expressing neurons may be part of an evolutionarily conserved pathway linking neurogenesis with inhibition of innate behavioral responses to stress.

To elucidate the mechanism through which Lef1 generates *crhbp*-expressing neurons, we have focused on the function of Lef1 target genes that encode developmental transcription factors known to regulate hypothalamic neurogenesis (Xie et al., 2017). We recently showed that one of these genes, *bsx*, is required for the differentiation of multiple Lef1-dependent hypothalamic subtypes including *crhbp*-expressing neurons (Schredelseker et al., 2020). Based on previous work showing that multiple transcription factors likely function in combination to promote the differentiation of specific hypothalamic neuron subtypes (Xie and Dorsky, 2017), we hypothesized that other Lef1 targets might also be required for *crhbp* expression, and evidence indicated that the *Orthopedia* paralogous gene *otpb* could play such a role. Some behavioral phenotypes reported in *otpb* mutants were consistent with the decreased exploration that we observed in *lef1* mutants using the novel tank diving assay (Xie et al., 2017). While only subtle developmental defects were reported in *otpb* mutant larvae (Wircer et al., 2017), *crhbp* expression was not assessed in these studies.

Here we show that expression of *otpb*, but not the paralogous gene *otpa*, is Lef1-dependent in the developing zebrafish PR. We find that expression of *crhbp* and *nos1* are absent in the PR of *otpb* mutants, while other Lef1-dependent genes are unaffected. Transgenic analysis of a conserved noncoding DNA element upstream of the *crhbp* promoter drives *lef1*-dependent, *otpb*-independent reporter expression in the PR, suggesting a transcriptional regulatory network of Lef1 targets. Finally, we show that *otpb* mutants exhibit decreased exploration in a novel tank diving assay, consistent with an increased stress response. Together, our data indicate a potential mechanism for the modulation of innate behavior by Lef1, through the regulation of *otpb*-dependent *crhbp* expression during hypothalamic neurogenesis.

## Results and Discussion

### *otpb* expression in the zebrafish PR is *lef1*-dependent

To fully characterize the expression of *otpb* and confirm that it is a Lef1 target in the developing zebrafish hypothalamus, we performed in situ hybridization on *lef1* mutant larvae at 3 days post-fertilization (dpf). We found that *otpb* was expressed throughout the hypothalamus, and that expression was decreased specifically in the lateral PR of homozygous *lef1* mutants (Xie et al., 2017) (Fig. 1A). Interestingly, this is the same region where *otpb* is expressed in the absence of *otpa* (Fernandes et al., 2013). We therefore repeated the experiment using a probe for *otpa*, and found that there was no obvious difference in expression between controls and *lef1* mutants (Fig. 1B), suggesting that *otpb* may have a unique Lef1-dependent function in the PR.

**Figure 1.**
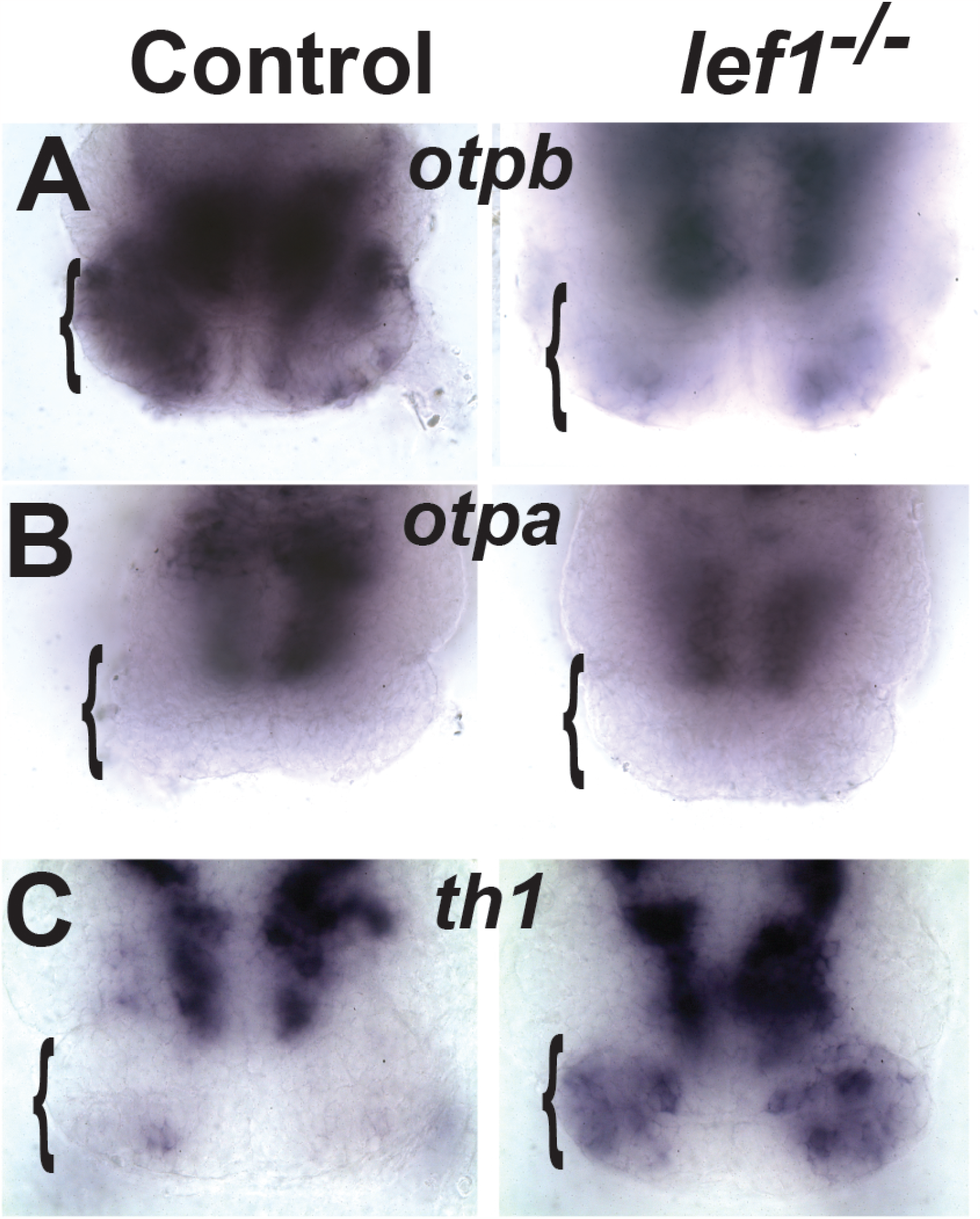
*lef1* is required for expression of *otpb*, but not *otpa*, in the PR. (A) In situ hybridization with a probe for *otpb* mRNA shows reduced expression in the PR of homozygous *lef1* null mutants compared to control siblings. (B) Analysis with a probe for *otpa* shows equivalent expression in the PR of homozygous *lef1* mutants and controls. (C) Analysis with a probe for *th1* shows ectopic expression in the lateral PR of homozygous *lef1* mutants. Representative ventral views of dissected brains at 3 dpf are shown, with brackets indicating the hypothalamic PR.

The absence of *otpb* expression in the PR of *lef1* mutants coincides with the loss of multiple neuron subtype markers (Xie et al., 2017), raising the question of whether the primary defect in these mutants is a general failure of neurogenesis, or instead a mis-specification of neural progenitors due to the function of Lef1-independent transcription factors. While screening *lef1* mutants for expression of other hypothalamic neuron subtype markers, we made the unexpected observation that *tyrosine hydroxylase 1* (*th1*) was ectopically expressed in lateral regions of the PR (Fig. 1C). Previous work reported that generation of *th1*-expressing dopaminergic neurons in the zebrafish diencephalon requires redundant functions of *otpa* and *otpb*, but is unaffected in single *otpb* mutants (Fernandes et al., 2013). These results suggest that the combined roles of *otpa* and *otpb* in generating dopaminergic neurons may be to antagonize Lef1 function in specifying a different neuron subtype.

### *otpb* function is required for expression of *crhbp* and *nos1* in the lateral PR

To determine whether *otpb* regulates *crhbp* expression in the hypothalamus, we performed *in situ* hybridization on *otpb*^*sa115*^ mutant larvae at 3 dpf. The *sa115* allele encodes a truncated protein lacking the homeodomain, and is thus likely to be a functional null mutation (Fernandes et al., 2013). We found reduced expression of *crhbp* in the lateral PR of homozygous *otpb*^*sa115*^ mutants compared to control siblings (Fig. 2A), coinciding with the loss of *otpb* expression in *lef1* homozygous mutants (Fig. 1A). In addition, we observed a similar reduction in the expression of another Lef1-dependent gene, *nos1* (Fig. 2B). This gene encodes an enzyme required for synthesis of the neuronal signal nitric oxide, and like *crhbp* is required for inhibiting innate stress response behavior (Carreno Gutierrez et al., 2020). The expression of both *crhbp* and *nos1* was only affected in the lateral PR, where *otpb* expression is Lef1-dependent and *otpa* is not expressed (Fig. 1). In contrast, the expression of *hdc*, another Lef1-dependent neuron subtype marker required for synthesis of the neuronal signal histamine, was unaffected in homozygous *otpb* mutants (Fig. 2C). We next tested 20 other Lef1-dependent genes encoding developmental transcription factors, neuron subtype markers, and known Wnt pathway targets (Xie et al., 2017), and found that they were all expressed normally in *otpb* mutants (data not shown). Together these results suggest that Otpb may function in combination with other Lef1 target transcription factors such as Bsx (Schredelseker et al., 2020), to generate a stress response-inhibiting *crhbp*+, *nos1*+ neuron subtype in the PR.

**Figure 2:**
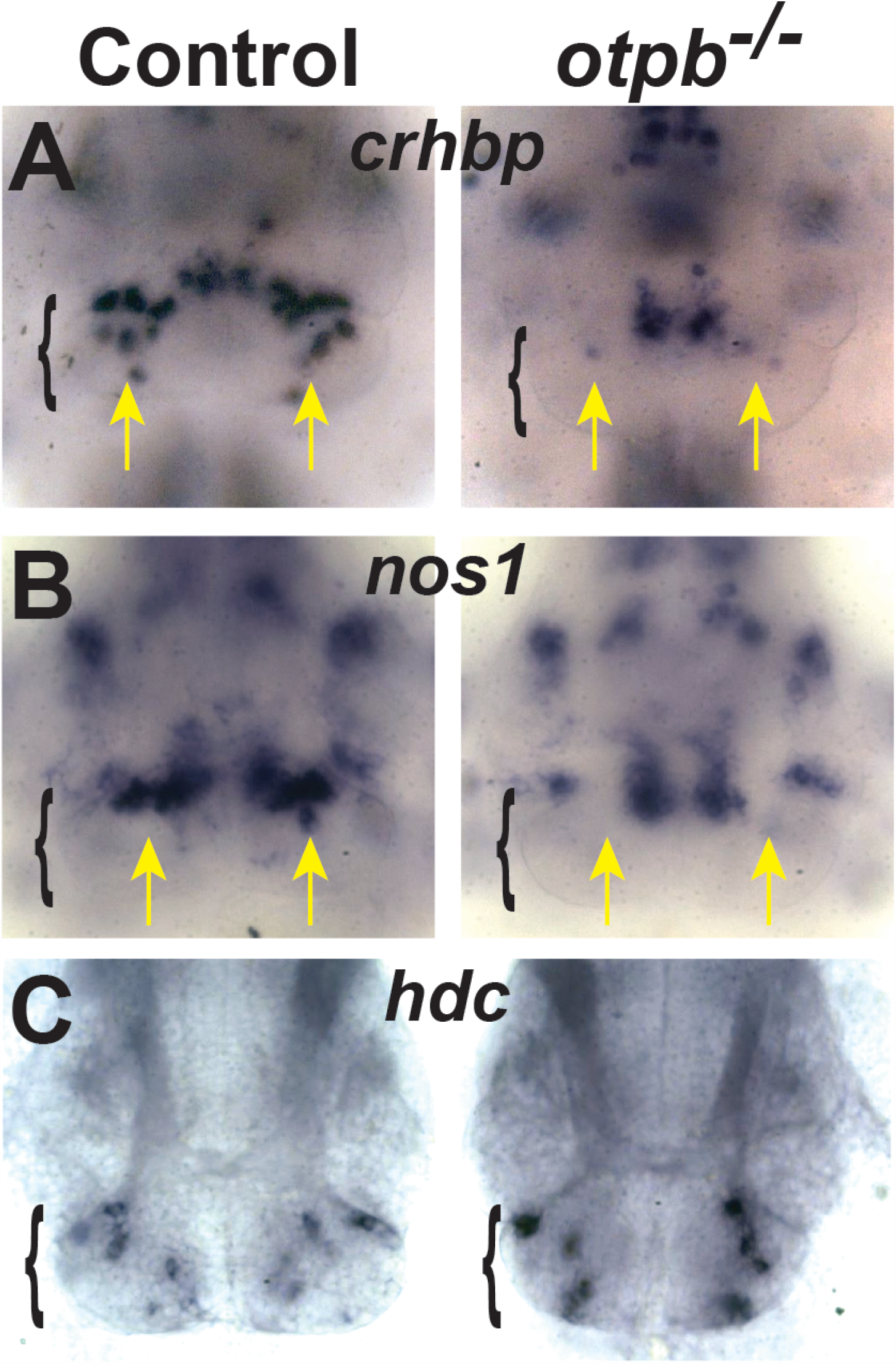
*otpb* is required for expression of *crhbp* and *nos1* in the PR. (A,B) In situ hybridization with probes for *crhbp* (A) and *nos1* (B) mRNA shows reduced expression specifically in the lateral PR of homozygous *otpb* null mutants (yellow arrows). (C) In contrast, *hdc* expression is unaffected in *otpb* null mutants. Representative ventral views of dissected brains at 3 dpf are shown, with brackets indicating the hypothalamic PR.

### *otpb* and *lef1* regulate *crhbp* expression in the PR through distinct transcriptional mechanisms

To understand the specific roles of *lef1* and *otpb* in regulating *crhbp* expression, we first used the UCSC genome browser (Kent et al., 2002) to identify predicted conserved noncoding elements upstream of the *crhbp* coding sequence. We then isolated a 381 bp fragment adjacent to the start codon that contained two regions of high conservation among fish species (Fig. 3A). The fragment also included a predicted transcription start site, and we used multisite Gateway cloning to assemble this fragment (*-381crhbp*) into a Tol2 GFP reporter plasmid construct. Injection of the plasmid with *tol2* mRNA into 1-cell embryos produced larvae with mosaic GFP expression in the PR at 3 dpf, which were then raised to adulthood. The progeny of several F0 founders exhibited GFP expression throughout the PR and in other brain regions (Fig. 3B), and these F1 larvae were raised and outcrossed. An F2 family with reproducible GFP expression that showed mendelian inheritance was used to establish a new transgenic allele: *-381crhbp:gfp*^*zd22Tg*^.

**Figure 3:**
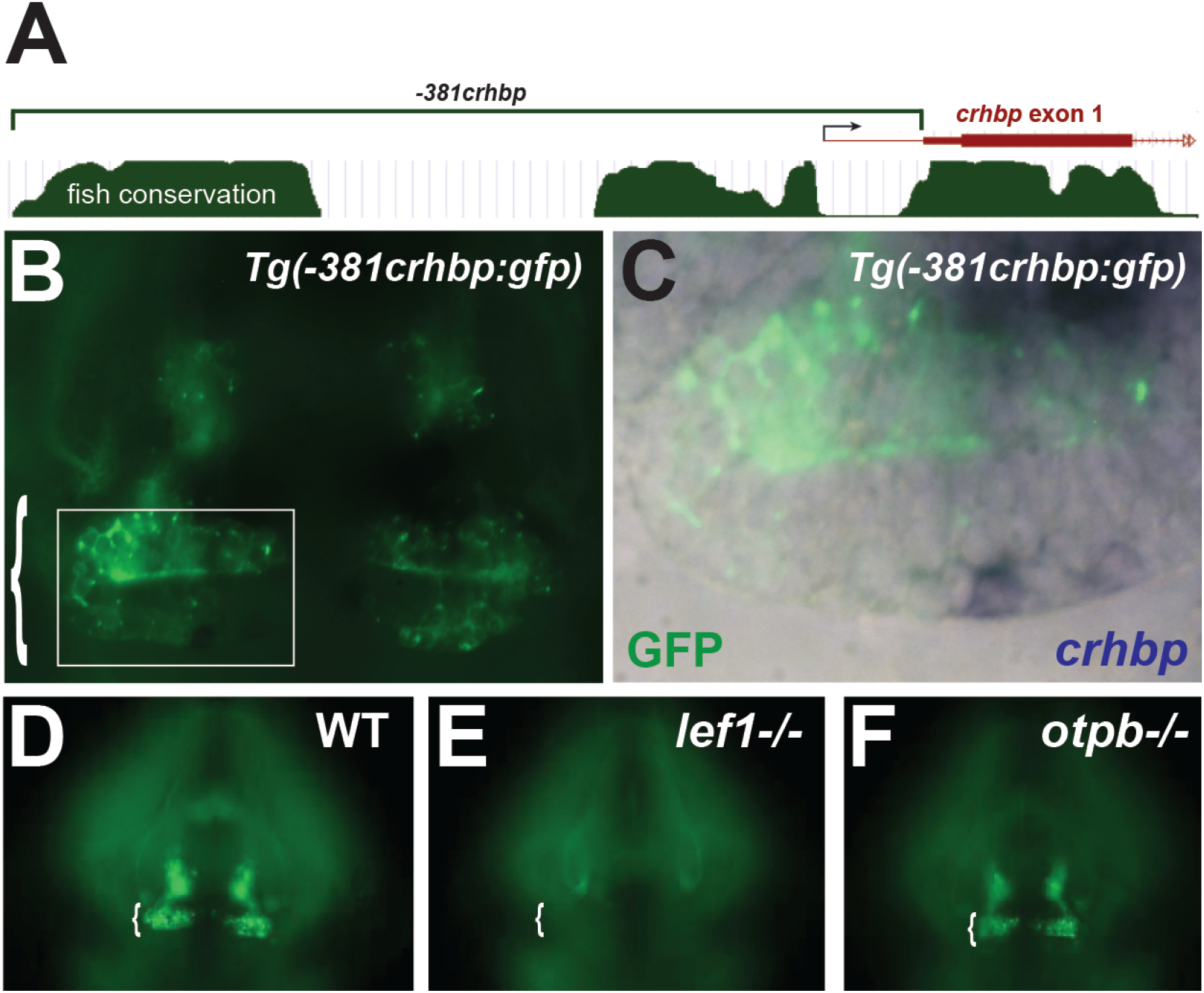
A conserved noncoding element upstream of *crhbp* drives reporter expression in the PR. (A) Schematic diagram of the *-381crhbp* noncoding element adapted from the UCSC Genome Browser. Sequence conservation among fish species is depicted in green and arrow indicates predicted transcription start site. (B) GFP expression in the PR of a *Tg(−381crhbp:gfp)* transgenic larva at 3 dpf, with area enlarged in panel C outlined by white box. (C) Simultaneous detection of GFP reporter expression (green) and *crhbp* mRNA (blue) indicates a lack of correlation within individual PR cells. (D-F) Comparison of GFP expression in the PR of wild-type (D), homozygous *lef1* mutant (E), and homozygous *otpb* mutant (F) larvae. Only GFP-labeled axons from other brain regions are visible in *lef1* mutants, while expression appears to be unaffected in the PR of *otpb* mutants. Representative ventral views of dissected brains at 3 dpf are shown, with brackets indicating the hypothalamic PR.

We next tested whether the reporter transgene co-localized with hypothalamic *crhbp* mRNA expression using combined GFP immunohistochemistry and in situ hybridization. We found that the level of reporter activity did not correlate with the level of *crhbp* mRNA, and that many GFP-positive cells in fact appeared to express low levels of *crhbp* mRNA (Fig. 3C). We then examined the regulation of reporter expression by *lef1*, and observed a complete absence of GFP expression in the PR of *lef1* mutants at 3 dpf, consistent with the loss of neurons observed in this region (Fig. 3D,E). However, we were surprised to find that GFP expression in the PR of *otpb* mutants was indistinguishable from controls (Fig. 3D,F). Together, these data indicate that Lef1 regulates *crhbp* expression in the lateral PR either directly or indirectly through this conserved noncoding element, and that Lef1-dependent regulation through *otpb* occurs through a distinct mechanism.

### *otpb* mutants exhibit decreased exploration of a novel environment

If the decreased exploration observed in *lef1* mutants is mediated by *crhbp*-expressing hypothalamic neurons, and *crhbp* expression in these neurons is lost in *otpb* mutants, we reasoned that both mutants might exhibit similar behavioral phenotypes. We first confirmed that *crhbp* expression in the PR was reduced in *otpb* mutants at 21 dpf (Fig. 4 A,B), and then assayed their behavior at this stage using a novel tank diving assay (Cachat et al., 2010). We found that like *lef1* mutants, *otpb* mutants exhibited significantly increased latency to enter the upper zone of the tank (Fig. 4C), and a significant decrease in total swimming distance (D) which may be partially due to a significant increase in time spent immobile (Fig. 4E).

**Figure 4:**
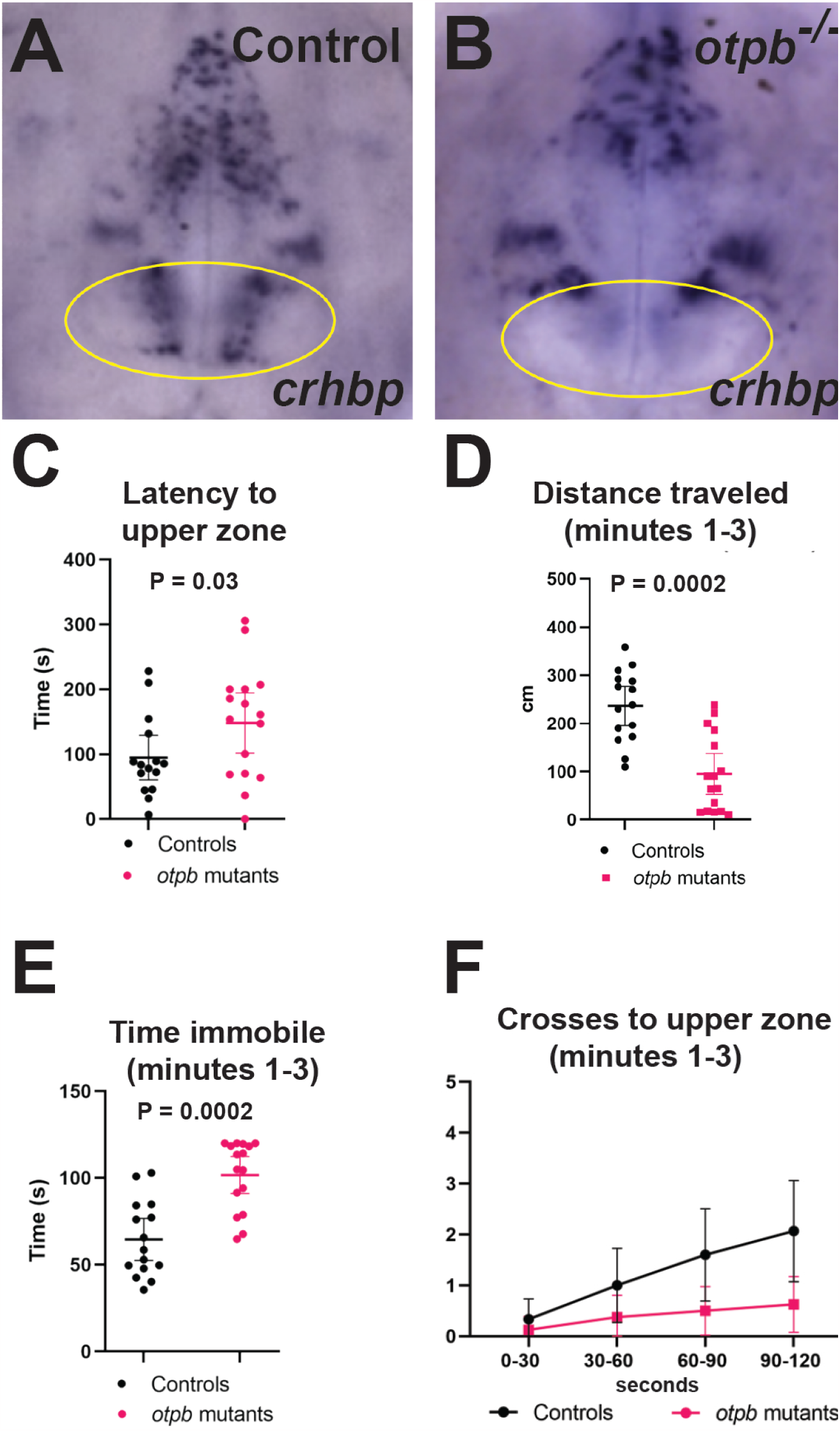
*otpb* mutants exhibit decreased exploratory behavior. (A,B) *crhbp* expression is reduced in the PR (yellow oval) of homozygous *otpb* mutants compared to wild-type or heterozygous control siblings. (C) *otpb* mutants are significantly delayed in exploration in the upper zone of a novel tank. (D,E) *otpb* mutants exhibit a significant decrease in total distance traveled (D) and spend significantly more time immobile (E) during minutes 1-3 compared to controls. (F) Two-way ANOVA analysis shows a significant decrease in crosses into the upper zone at each 30 second timepoint, and a significant difference in overall acclimation rate between *otpb* mutants and controls during minutes 1-3 in a novel tank. N=15 controls and 18 homozygous mutants for (C-E), N=15 controls and 15 homozygous mutants for (F). Error bars in all graphs indicate 95% CI.

To compare rates of acclimation to the novel environment, we measured crosses to the tank upper zone in four 30 second intervals during the first 1-3 minutes (Fig. 4F). While all larvae significantly increased their number of crosses to the upper zone during the 1-3-minute interval, controls had a faster acclimation rate than *otpb* mutants. These results are all consistent with the loss of *crhbp* expression in both *lef1* and *otpb* mutants, based on the known function of Crhbp in inhibiting the Crh-mediated stress response in other species.

## Conclusions

The data presented here support a function for *otpb* in a Lef1-dependent transcriptional network required for expression of *crhbp* in PR neurons that inhibit stress response behavior in zebrafish. While individual Lef1 target genes and Lef1-dependent neuron subtypes can vary between species, our results support a model in which inhibiting the innate behavioral response to stress may be an evolutionarily conserved role. While the specific neuronal functions and behavioral circuits underlying this role remain unclear, they likely include a circuit that is modulated by the release of neuropeptides and hormones. Such molecules remain to be identified and/or implicated in a specific Lef1-dependent behavioral response.

The duplication-degeneration-complementation (DDC) evolutionary model (Force et al., 1999) proposes that duplicate paralogous genes are retained through complimentary degenerative mutations in regulatory elements that partition ancestral functions. This model therefore suggests that the regulation of *otpb* by Lef1 may have been inherited from an ancestral *Otp* gene, and lost in zebrafish *otpa*. Based on the DDC model, it is possible that gene duplication has also revealed a Lef1-dependent role in stress response inhibition that was masked by other functions in species with single *Otp* orthologs.

### Experimental Procedures

#### Use of animals and establishment of *-381crhbp:gfp* transgenic line

All fish were used and handled in accordance with University of Utah IACUC guidelines. Wild-type AB strain, *lef1*^*zd11*^ (Wang et al., 2012), and *otpb*^*sa115*^ (Fernandes et al., 2013) zebrafish were crossed to generate embryos. PCR amplification for genotyping mutants was performed on fin clip DNA. To make the *Tg(−381crhbp:gfp)*^*zd22*^ allele, DNA injections were performed at the 1-cell stage. Injected F0 fish were outcrossed and embryos were visually inspected for the expression of GFP in the hypothalamus as well as a *cmlc2:egfp* transgene marker expressed in the heart.

#### Construction of *-381crhbp:gfp* transgene

Genomic sequences were PCR amplified from AB fish using primers that included restriction enzyme sites: Forward primer (FseI): TAGTCTACCAATTTGCAGTCAAATTTGTT Reverse primer (AscI):CCCGGCGCGCCGTCCGTTCAGTCTGTCCGCC Amplicons were digested with restriction enzymes, then ligated into a 5’ Gateway entry clone vector.

A Gateway LR reaction was then performed with the 5’ entry clone, a middle entry clone containing the *gfp* cDNA, a 3’ entry clone containing the SV40 polyA sequence, and a Tol2 destination vector to create a transgenic reporter plasmid.

#### In Situ Hybridization and Immunohistochemistry

In situ hybridization was carried out as previously described (Xie et al., 2017). PCR primers used to generate mRNA probe templates were:

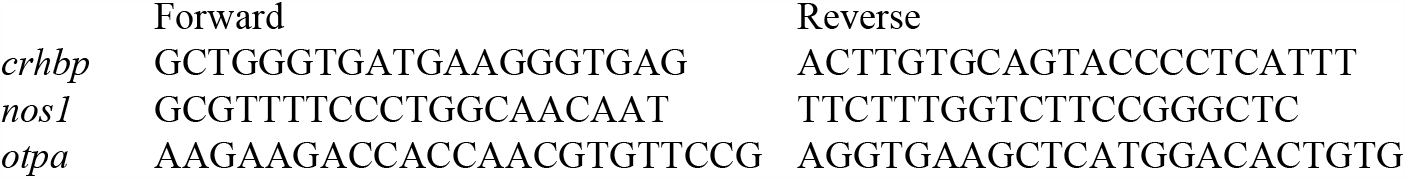
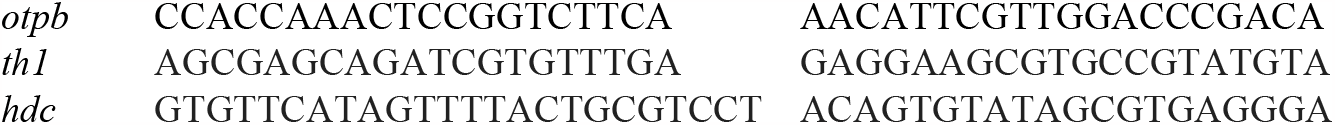

For simultaneous detection of GFP protein, brain digestion was performed with 0.1% collagenase instead of proteinase K, and a rabbit anti-GFP antibody was combined with the anti-digoxigenin antibody. Brains were then incubated in goat anti-rabbit Alexa 488 antibody before imaging.

#### Imaging and analysis

Whole mount imaging of embryos was performed on an Olympus SZX16 dissecting microscope, with fluorescence used to validate GFP expression in the *zd21* allele. Live embryos were imaged at 4X-20X magnification on an Olympus BX51WI fluorescent compound microscope at 3 dpf to document GFP expression patterns.

#### Novel tank diving test

Homozygous *otpb* mutant fish and control siblings were generated from *otpb*^*sa115*^ heterozygous mutant incrosses and raised in separate tanks. The novel tank diving test was performed on individual 21 dpf larvae in tanks illuminated with white light centered above and surrounded by white walls to avoid social interaction. Videos were acquired by a Logitech C920 HD Pro Webcam Full HD 1080p camera. One homozygous mutant and one control larva were assayed simultaneously, in alternating tank positions for each trial. Videos were analyzed using Ethovision XT version 11.5 (Noldus, Leesburg, VA) with a tracking period of 6 minutes, beginning 30 seconds after release. Tracks were analyzed for latency to initial cross into the upper zone, total distance travelled. time spent immobile, and number of crosses into the upper zone (Cachat et al., 2010).

#### Statistical Analysis

For measuring latency, distance traveled, and time immobile, a Student’s t-test was used to compare control and mutant larvae. For measuring crosses to the upper zone over time, a two-way ANOVA was used to compare three variables: 1) genotype, 2) behavior over time (acclimation rate), and 3) acclimation rates between genotypes.

